# Altered brain morphology in boys with attention-deficit/hyperactivity disorder with and without comorbid conduct disorder/oppositional defiant disorder

**DOI:** 10.1101/553552

**Authors:** Nora C. Vetter, Lea L. Backhausen, Judith Buse, Veit Roessner, Michael N. Smolka

**Author notes:** Correspondence to Nora C. Vetter, Universitätsklinikum Carl Gustav Carus, an der Technischen Universität Dresden, Fetscherstraße 74, 01307 Dresden, Germany. Shared senior-authorship (Veit Roessner, Michael N. Smolka).

## Abstract

About 50% of ADHD patients suffer from comorbidity with oppositional defiant disorder/conduct disorder (ODD/CD). Most previous studies on structural morphology did not differentiate between pure ADHD (ADHD-only) and ADHD with comorbid ODD/CD (ADHD+ODD/CD) and only focused on specific measures (e.g. volumetric differences), leading to inconsistent results. Therefore, we aimed to investigate the structural profile of ADHD-only versus ADHD+ODD/CD spanning different indices, i.e. subcortical and cortical volume, cortical thickness, and surface area. We aimed to disentangle disorder-specific etiological contributions regarding structural brain alterations and expected similar and differential alterations between the patient groups.

We acquired structural images from an adolescent sample range (11 to 17 years) matched with regard to age, pubertal status, and IQ - including 36 boys with ADHD-only, 26 boys with ADHD+ODD/CD, and 30 typically developing boys (TD). We analyzed subcortical and cortical volume, cortical thickness, and surface area with FreeSurfer.

We found reductions in total gray matter and right cerebellar volume as well as total surface area for both patient groups. For the left cerebellar volume ADHD+ODD/CD, but not ADHD only differed from TD. Boys with ADHD+ODD/CD had a thicker cortex than the other groups in a right rostral middle frontal cluster, which was related to stronger ODD/CD symptoms, even when controlling for ADHD symptoms. No group differences in local cortical volume or surface area emerged.

Overall, we found similarities but also differences in brain morphology between the two related disorders. Patients with a “double burden” seem to be even more affected than patients with pure ADHD.

## Introduction

One of the most commonly diagnosed childhood psychiatric disorders affecting 3 to 9% of children and adolescents is attention-deficit hyperactivity disorder (ADHD; Barkley, 2014). ADHD is defined by developmentally inappropriate levels of inattention, and/or hyperactivity/impulsivity (American Psychiatric Association, 1994). ADHD is related to academic and work-related difficulties and represents a major public health burden (Pardini & Fite, 2010).

Around 48 to 67 % of ADHD patients suffer from comorbid oppositional defiant disorder (ODD) or conduct disorder (CD(Connor, Steeber, & McBurnett, 2010). ODD is defined by developmentally inappropriate, negativistic, defiant, and disobedient behavior and is often a precursor for the more severe CD (Maughan, Rowe, Messer, Goodman, & Meltzer, 2004), which is characterized by aggression, deceitfulness, property destruction and rules violation. Compared with patients with pure ADHD, patients with ADHD and comorbid ODD/CD show an earlier age of onset for ADHD symptoms, and have a worse prognosis (Connor & Doerfler, 2008; Loeber, Burke, Lahey, Winters, & Zebra, 2000).

ADHD patients show neuropsychological deficits in attention and “cool” executive function such as inhibition and switching (Halperin & Schulz, 2006), while patients with ODD/CD seem to have difficulties in “hot” executive functions including reward and emotion processing (Byrd, Loeber, & Pardini, 2014). In line with these neuropsychological findings, morphological neuroimaging studies have shown abnormalities for ADHD in frontostriatal and frontocerebellar circuits and for ODD/CD in frontolimbic circuits (for a review see Rubia, 2011).

In detail, volume reductions for ADHD have been most consistently reported for the striatum, the cerebellum, and total gray matter (GM; Greven et al., 2015; for metaanalyses see Hoogman et al., 2017; Nakao, Radua, Rubia, & Mataix-Cols, 2011; Valera, Faraone, Murray, & Seidman, 2007). Regarding frontal alterations, studies reported a decreased (Ambrosino, de Zeeuw, Wierenga, van Dijk, & Durston, 2017) or increased (Garrett et al., 2008; Semrud-Clikeman, Pliszka, Bledsoe, & Lancaster, 2014) volume of the prefrontal cortex.

Some studies also found a reduction in prefrontal cortical thickness (CT; Almeida et al., 2010; Hoekzema et al., 2012; Shaw et al., 2006; Yang, Carrey, Bernier, & MacMaster, 2015) or a delayed cortical thinning (Shaw et al., 2007). In contrast, further research shows an increased CT in other areas (occipital or temporal cortex, pre-supplementary motor area; Almeida Montes et al., 2013; Duerden, Tannock, & Dockstader, 2012) or no CT alterations (Ambrosino et al., 2017; Wolosin, Richardson, Hennessey, Denckla, & Mostofsky, 2009).

The few recent studies to investigate an additional dimension of cortical morphology, namely surface area (SA) have reported reduced total, frontal, temporal or parietal SA (Dirlikov et al., 2015; Wolosin et al., 2009). Taken together, while overall frontostriatal and frontocerebellar circuits seem to be affected in ADHD, previous studies partly remain inconsistent, mainly for the frontal alterations (Stevens & Haney-Caron, 2012).

For ODD/CD, volume reductions were found in the striatum, amygdala, hippocampus, or prefrontal cortex (Noordermeer, Luman, & Oosterlaan, 2016; Rogers & Brito, 2016). Reports on prefrontal volume also include increases (De Brito et al., 2009). Very few ODD/CD studies investigated CT and SA. These reported CT reductions for temporal and parietal (Hyatt, Haney-Caron, & Stevens, 2012; Wallace et al., 2014) or prefrontal regions (Fahim et al., 2011). Prefrontal SA reductions (Fairchild et al., 2015; Sarkar et al., 2015) or no SA differences (Wallace et al., 2014) have been also been shown. Summarizing these findings, while ODD/CD alterations are mainly evident in frontolimbic circuits, several inconsistencies are evident regarding the specific frontal alterations.

Divergent findings for both ADHD and ODD/CD studies might stem from previous samples that partly included ADHD patients comorbid with ODD/CD (Wolosin et al., 2009) or vice versa (see Stevens & Haney-Caron, 2012 for a summary). Thus, the contribution of the respective other disorder could only – if at all – be statistically controlled for. Studies on ADHD+ODD/CD that try to disentangle each disorder’s specific contribution are rare. One study contrasted adolescent ADHD vs. CD volume abnormalities and found total GM volume reductions reflecting frontal, temporal, parietal, and subcortical volume reductions in CD but not ADHD patients (Stevens & Haney-Caron, 2012). The authors therefore questioned previously reported frontal abnormalities in ADHD and speculated they might stem from ODD/CD comorbidity in previous ADHD samples. To date, only one recent study compared structural morphology (volume and thickness) in adolescents with pure ADHD to those with ADHD+ODD and found both similar and differential cortical volume (CV) alterations (Noordermeer et al., 2017). Both patient groups showed alterations from TD in the prefrontal cortex. Patients with ADHD+ODD had a stronger reduction in the orbito-, middle- and superior frontal cortex, which is in line with stronger impairments in executive functions (Connor & Doerfler, 2008; Loeber et al., 2000). Differential alterations were found for ADHD+ODD differing from pure ADHD and TD in some prefrontal regions of interest (ROIs) and the precuneus and for ADHD+ODD differing from TD in the left middle temporal gyrus. No thickness or subcortical volume alterations emerged, possibly due to the rather old sample (mean age 16-17 years) and wide age range (7-29 years). Overall, inconsistencies in previous ADHD studies might stem from large age differences ranging from child (Dirlikov et al., 2015; Wolosin et al., 2009; Yang et al., 2015) to adolescent (Almeida Montes et al., 2013; Ambrosino et al., 2017; Duerden et al., 2012) or adult samples (Fairchild et al., 2015) or samples spanning all three age groups (Greven et al., 2015). Investigations on adolescent ADHD are rare (for a review see Lin & Roth, 2017). Brain development, specifically of the frontal cortex, continues well into adolescence (Gogtay et al., 2004). Therefore, fostering knowledge on the structural alterations of ADHD during this age phase by assessing adolescent samples with a more narrow age range would shed light on the neurodevelopment of this disorder. Overall, more investigations on the structural abnormalities associated with ADHD+ODD/CD compared to pure ADHD are warranted to reveal similarities and differences in the morphological alterations of the disorders. This might have important implications for disorder etiology and treatment.

Therefore, our major aim was to provide knowledge to disentangle morphological characteristics of adolescents with pure ADHD (ADHD-only) and ADHD+ODD/CD. Few previous studies analyzed several cortical and subcortical morphological markers simultaneously to reach an overall picture on alterations and no study did so for the comorbidity of ODD/CD. We therefore aimed at investigating the structural profile of both ADHD-only and ADHD+ODD/CD spanning (sub)cortical volume, CT, and SA. We assessed a well matched sample spanning the adolescent age phase (11 to 17 years). We included only boys due to the larger prevalence for males in ADHD (Willcutt, 2012) and the gender-specific morphology in ADHD (Seymour et al., 2017)

First, we predicted a reduced total GM volume in patient groups similar to Noordermeer et al. (2017). Second, for subcortical volumes, we predicted a reduced striatum and cerebellum (see megaanalysis by Hoogman et al., 2017) with a stronger reduction in ADHD+ODD/CD due to stronger symptoms and neuropsychological deficits observed in this disorder. In line with previous studies on ODD/CD (Rogers & Brito, 2016) we predicted for ADHD+ODD/CD a reduced volume of the amygdala and hippocampus.

We additionally explored, on a whole-brain basis, cortical alterations in volume, thickness, and SA. We expected ADHD+ODD/CD to be a hybrid form of ADHD-only and pure ODD/CD (Schachar & Tannock, 1995) and thus yield more severe alterations than each disorder alone (McAlonan et al., 2007). We expected such alterations mainly in the prefrontal cortex (see also Noordermeer et al., 2017 and Stevens & Haney-Caron, 2012)

## Methods

### Participants and study design

This study was part of larger neuroimaging study on ADHD and comorbid ODD/CD using another subsample than our previous publications (Backhausen et al., 2016; Vetter et al., 2018). It was carried out according to the latest version of the Declaration of Helsinki and approved by the ethics committee of the TU Dresden. Participants and parents or legal guardians gave their written informed consent and participants received around 40 €.

We recruited three groups of boys within the age range of 11-17 years: First, ADHD-only with 51 boys diagnosed only with ADHD; second, ADHD+ODD/CDADHD+ODD/CD with 29 boys diagnosed with ADHD and comorbid ODD/CD, and third TD with 40 boys without any psychiatric diagnosis. Exclusion criteria consisted of IQ<80 or any other additional axis-I disorder. We recruited patients within the local in- and outpatient clinics and TD boys in schools, doctors’ offices and a parish. Diagnoses were made according to the ICD-10 (World Health Organization, 1992) by board certified child and adolescent psychiatrists and verified by the Mini International Neuropsychiatric Interview for Children and Adolescents (Sheehan et al., 1998).

We had to exclude nine boys due to motion (three ADHD-only, one ADHD+ODD/CD, two TD), ferromagnetic artifacts (one ADHD-only), or gross neuroanatomical abnormalities (one ADHD-only, one TD). The remaining sample (n=46 ADHD-only, n=28 ADHD+ODD/CD, and n=37 TD) was not matched regarding age, IQ, socioeconomic or pubertal status. To achieve a matched group at least for the most important variables (age, pubertal status, and IQ), we had to exclude the youngest ADHD+ODD/CD patients (n=2), and those ADHD-only patients with a very low (n=10) and TD with a very high IQ (n=7). This resulted in a sample of n=36 ADHD-only, n=26 ADHD+ODD/CD and n=30 TD (Table 1).

**Table 1.** Demographics, clinical characteristics and group comparisons (n = 92)

| Parameter |  | ADHD-only (N=36) | ADHD+ODD/CD (N=26) | TD (N=30) | Group comparison |  |  | Post-Hoc |
| --- | --- | --- | --- | --- | --- | --- | --- | --- |
|  |  | M (SD) | M (SD) | M (SD) | F | df | P |  |
| Age in years |  | 13.02 (1.57) | 12.99 (1.17) | 13.58 (1.58) | 1.46 | 2, 89 | n.s. |  |
| Socioeconomic status<sup>a</sup> |  | 12.41 (3.83) | 9.79 (4.07) | 15.65 (3.82) | 13.67 | 2, 73 | < .001 | ADHD-only<TD<sup>1</sup>,<br>ADHD+ODD/CD<TD<sup>1</sup> |
| IQ<sup>b</sup> |  | 107 (8) | 106 (13) | 110 (8) | 1.46 | 2, 89 | n.s. |  |
| Pubertal status<sup>c</sup> |  | 2.45 (.94) | 2.13 (.82) | 2.68 (1) | 1.4 | 2, 81 | n.s. |  |
| CBCL<sup>d</sup> | Attention problems | 66.13 (7.016) | 70.61(8.68) | 54.41(5.71) | 35.26 | 2, 77 | <.001 | ADHD-only>TD<sup>1</sup>,<br>ADHD+ODD/CD>TD<sup>1</sup> |
|  | Rule-breaking behavior | 56.10 (6.71) | 66.52 (7.43) | 52.93 (4.34) | 31.64 | 2, 77 | <.001 | ADHD-only>TD<sup>1</sup>.<br>ADHD+ODD/CD>TD<sup>1</sup> |
|  | Aggressive behavior | 60.93 (9.68) | 73.83 (9.43) | 54.48(6.39) | 31.97 | 2, 77 | <.001 | ADHD-only>TD<sup>2</sup>,<br>ADHD+ODD/CD>TD<sup>1</sup>,<br>ADHD-only<ADHD+ODD/CD<sup>1</sup> |
|  | Total | 61.37 (7.25) | 72.04 (6.5) | 51.44 (9.46) | 42.49 | 2, 77 | <.001 | ADHD-only>TD<sup>1</sup>,<br>ADHD+ODD/CD>TD<sup>1</sup>,<br>ADHD-only<ADHD+ODD/CD<sup>1</sup> |

*Note:* *F* values and *p* level refer to the main effect of group in ANOVA. <sup>1</sup>*p*<.01. <sup>2</sup>*p*<.05. Post-hoc tests are t-tests. ADHD-only= attention-deficit hyperactivity disorder group, ADHD+ODD/CD = comorbid attention-deficit hyperactivity disorder and conduct disorder/oppositional defiant disorder group, TD=typically developing group.
<sup>a</sup> Calculation of socioeconomic status included parents' school education. professional education. recent professional status and family income following the procedure suggested by Winkler and Stolzenberg (2009). Scores for mothers and fathers were averaged into a family-based measure of socioeconomic background. The score ranges from 3 to 21 with higher values indicating higher socioeconomic status.
<sup>b</sup> To estimate general cognitive ability the subtests Vocabulary, Letter-Number Sequencing, Matrix Reasoning and Symbol Search from the Wechsler Intelligence Scale For Children [WISC-IV. German adaptation; (Petermann & Petermann, 2010)] were used.
<sup>c</sup> Pubertal status ranges from 1 for prepubertal to 5 for postpubertal status, measured with the Pubertal Development Scale (Petersen, Crockett, Richards, & Boxer, 1988).
<sup>d</sup> CBCL – Child Behavior Checklist (Achenbach, 1991). Reported are CBCL *t* scores.
Please note that due to assessment difficulties some values are missing for a few participants (n range from 92 of 92 to 80 of 92).

Nine ADHD-only and four ADHD+ODD/CD patients had no medication history. Seventeen ADHD-only and ten ADHD+ODD/CD patients were taking methylphenidate, but paused medication 48 hours before scanning. Six ADHD+ODD/CD patients were simultaneously taking methylphenidate and atypical antipsychotics. Three ADHD-only patients had a history of other medication.

## Structural imaging

### Image acquisition and processing

We acquired magnetic resonance imaging scans on a 3T whole-body MR tomograph (Magnetom TRIO, Siemens, Erlangen, Germany) equipped with a 12-channel head coil and processed T1-weighted images using the cross-sectional pipeline of the segmentation and image analysis software FreeSurfer (Version 5.1, https://surfer.nmr.mgh.harvard.edu/). We applied detailed visual quality control for raw and processed images (Backhausen et al., 2016). See Supplement S1 for further details.

### Statistical analysis

Statistical analyses on demographic and clinical measures were performed with SPSS (IBM SPSS Statistics for Windows, Version 25.0, Armonk, NY, USA) using analyses of variance (ANOVAS). Using FreeSurfer’s volume-based stream we analyzed global measures (including intracranial volume, (ICV), total, cortical, and subcortical GM volume, mean CT, and total SA) and volume-based ROIs (including the cerebellum, thalamus, putamen, pallidum, hippocampus, amygdala, nucleus accumbens, and caudate). We ran ANOVAs followed by post-hoc t-tests to compare groups.

As cortical measures, we analyzed CV, SA and CT using FreeSurfer’s surface-based stream applying a whole brain approach with general linear modeling. First, to enable comparability with previous publications, we reported clusters surviving a cluster-forming threshold (CFT) of *p*<.05 with a precomputed Monte Carlo Z simulation with 10,000 iterations and a full-width at half-maximum Gaussian smoothing kernel of 15 mm. Then we performed robustness analyses. First, we applied new stringent thresholds (Greve & Fischl, 2018): CFT *p*<.005 for CT and CV, and *p*<.001 for SA with a cluster-wise threshold (CWP) of *p*<.05 for CV, CT, and SA. Second, for CV and SA we added ICV as a covariate (Greve & Fischl, 2018).

## Results

### Global volume, thickness, and surface area

We found 4 – 6% decreases for both patient groups compared to TD in the ICV (5% ADHD-only versus TD: *p*=.011, 5% ADHD+ODD/CD versus TD: *p*=.0013), total GM volume (4% ADHD-only versus TD: *p*=.013, 5% ADHD+ODD/CD versus TD: *p*=.009), right cortical GM volume (4% ADHD-only versus TD: *p*=.034, 5% ADHD+ODD/CD versus TD: *p*=.0035), and total left and right SA (left: 5% ADHD-only versus TD: *p*=.014, 6% ADHD+ODD/CD versus TD: *p*=.007; right: 4% ADHD-only versus TD: *p*=.024, 6% ADHD+ODD/CD versus TD: *p*=.006). Patient groups did not differ from each other. There were no group differences for subcortical GM volume or mean CT (see Table S1, available online).

### Volume-based ROIs

After including ICV as a covariate, cerebellar volume decreases of 5 to 6% remained significant (right: ADHD-only versus TD: *p*=.041, ADHD+ODD/CD versus TD: *p*=.005; left: ADHD+ODD/CD versus TD, *p*=.033; Table S1; Figure 1). See Table S1 (available online) for uncorrected results. After applying FDR correction, no group difference remained significant.

**Figure 1:**
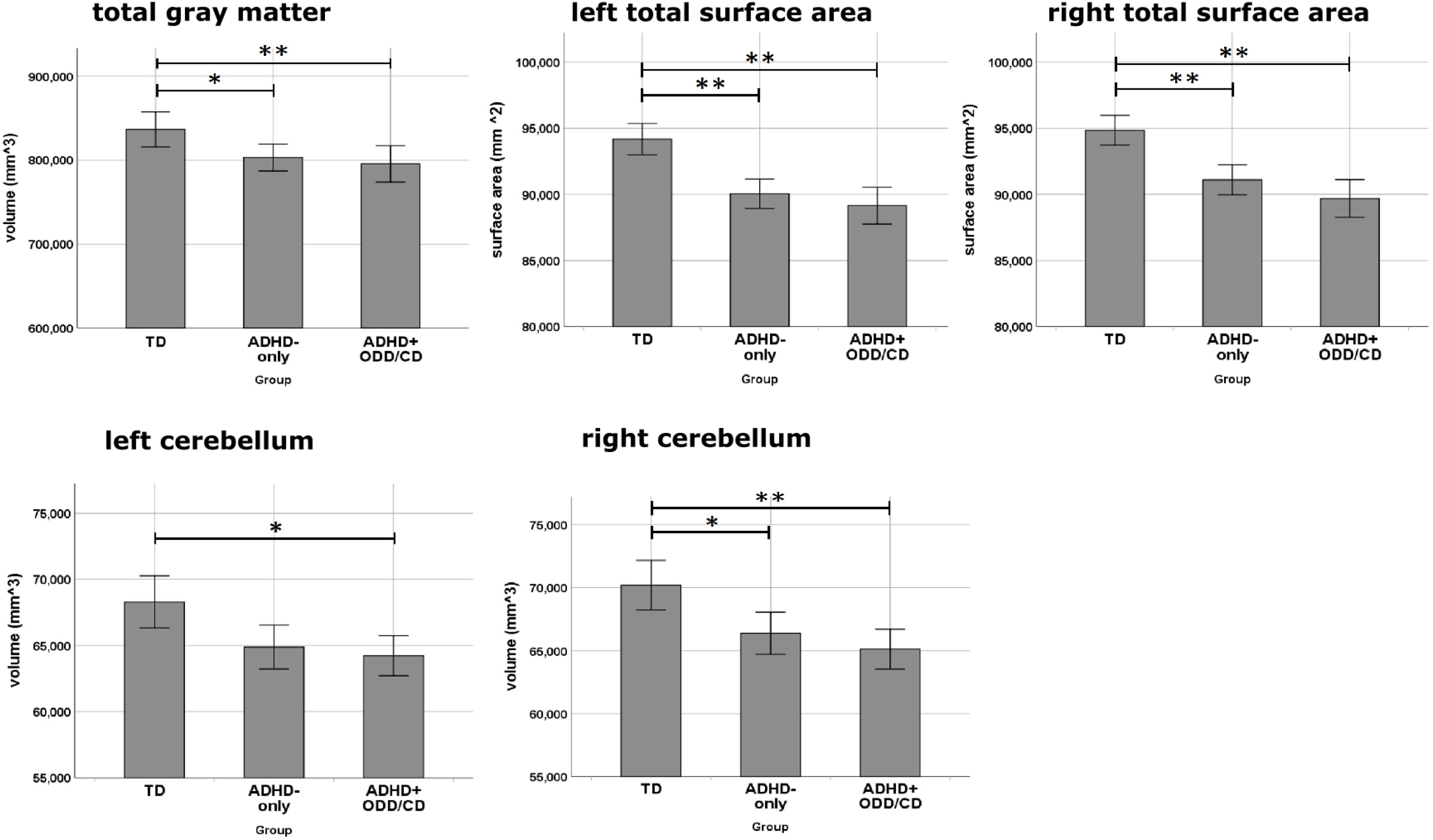
Global and subcortical volume differences. Note: \*\**p*<.01. \**p*<.05. Error bars denote standard errors. ADHD-only=group of boys diagnosed with ADHD only. ADHD+ODD/CD=group of boys diagnosed with ADHD and comorbid ODD/CD. TD= group of typically developing boys.

### Cortical thickness

We report anatomical regions according to the Desikan-Kiliany atlas (Desikan et al., 2006). For a liberal threshold (CFT *p*<.05), a main effect of group in a cluster of the right rostral middle frontal cortex emerged (see Table S2 for post-hoc tests, available online). When applying new stringent thresholds with a CFT *p*<.005 and CWP of *p*<.05 no cluster emerged for the main effect of group (Table 2). Since we had a priori hypotheses about differences between ADHD-only and TD as well as between ADHD-only and ADHD+ODD/CD, we performed post-hoc analyses. These revealed a 6.5 % increased CT for ADHD+ODD/CD versus TD and 7% increase for ADHD+ODD/CD versus ADHD-only in a rostral middle frontal cluster (Table 2, Figure 2).

**Table 2.**
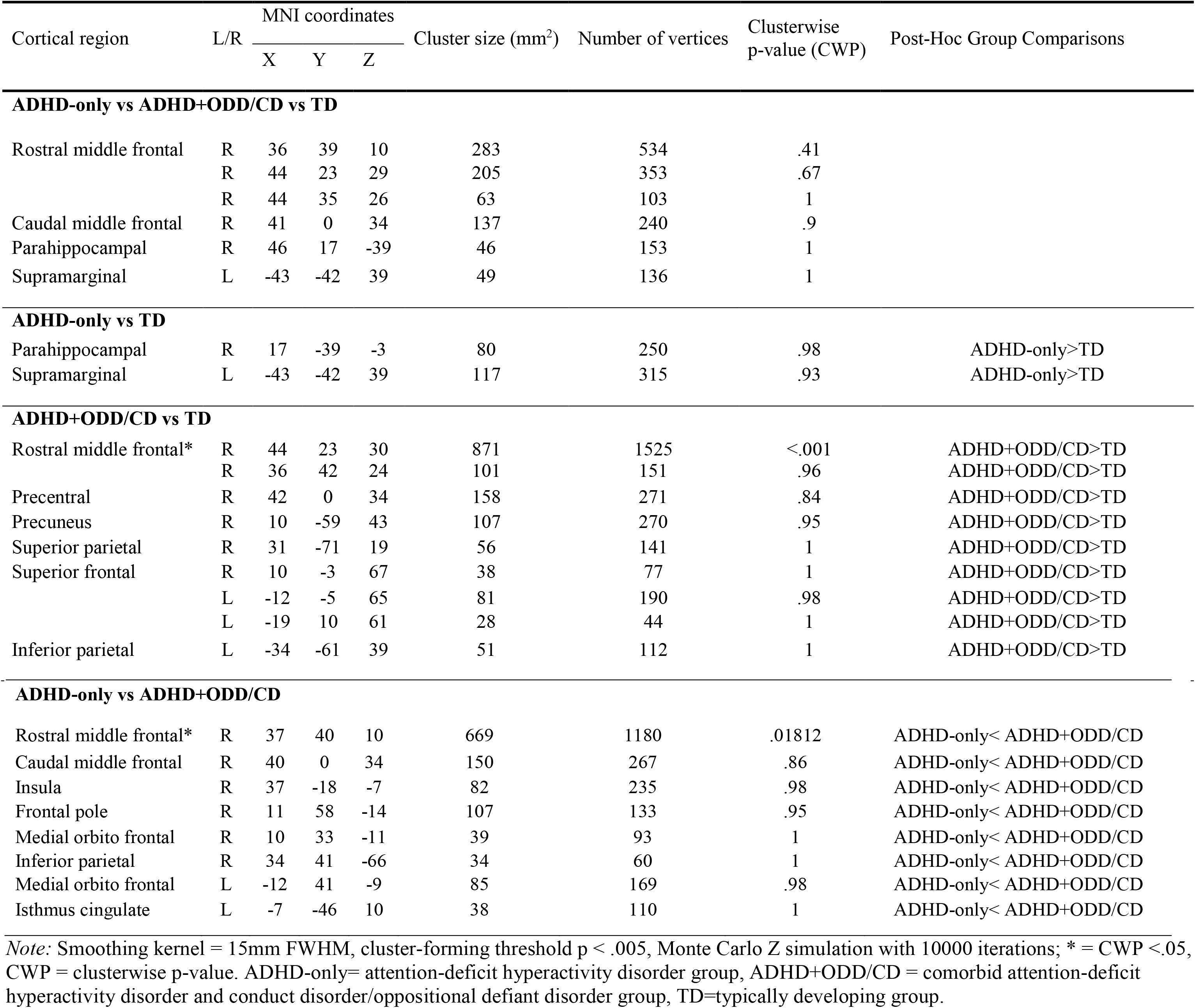
Differences in cortical thickness

**Figure 2:**
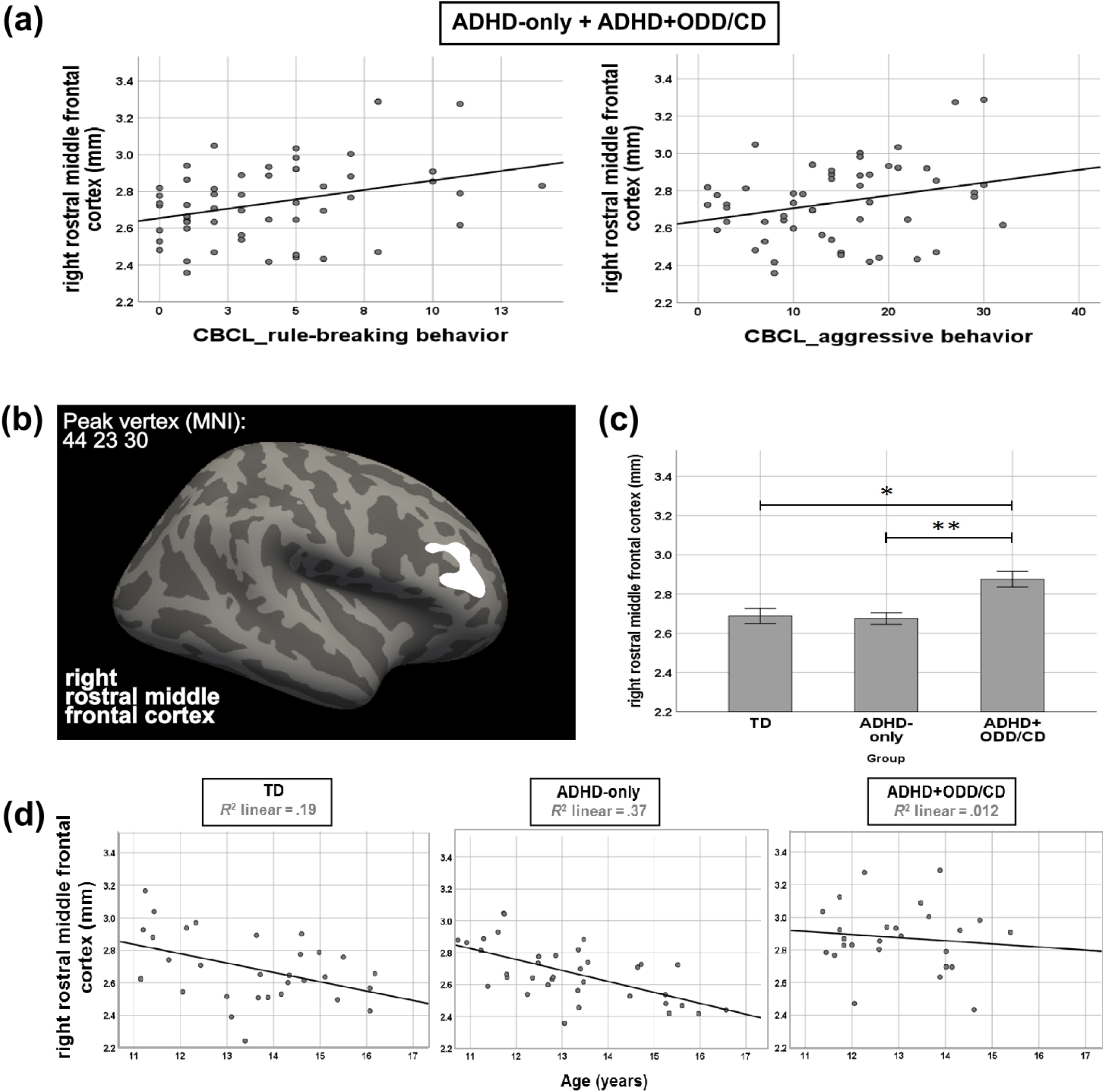
Cluster with group differences in cortical thickness. Note: (a) For contrasting ADHD-only versus ADHD+ODD/CD the only significant cluster is depicted (cluster forming threshold *p*<.005 and clusterwise *p*<.05) on the lateral surface of an inflated brain. (b) Mean cortical thickness for TD, ADHD-only and ADHD+ODD/CD extracted from the cluster shown in (a). Error bars denote standard errors. (c) Associations of mean cortical thickness in the cluster shown in (a) with raw scores of the rule-breaking and aggressive behavior scales as assessed with the CBCL. (d) Correlations with age of mean cortical thickness in the cluster shown in (a). \*\**p*<.01. \**p*<.05. ADHD-only=group of boys diagnosed with ADHD only. ADHD+ODD/CD=group of boys diagnosed with ADHD and comorbid ODD/CD. TD= group of typically developing boys

### Cortical volume and surface area

See Table S2 (available online) for results with a liberal threshold (CFT *p*<.05) and Table S3 (available online) for a stingent threshold (for CV: CFT *p*<.005; for SA: CFT *p*<.001). These results did not survive a CWP of *p*<.05 nor robustness analyses when including ICV as a covariate.

### Correlations of cortical thickness in the right rostral middle frontal cortex with symptom severity and age

To disentangle the contribution of disorder-specific symptoms to altered CT we correlated in both patient groups mean CT from the right rostral middle frontal cluster resulting for ADHD+ODD/CD versus ADHD-only with ADHD symptoms (scale “attention problems”) controlling for ODD/CD symptoms (scale “externalizing behavior” consisting of “aggressive” and “rule-breaking behavior”) and with ODD/CD symptoms controlling for ADHD symptoms as assessed with the CBCL.

CT correlated with ODD/CD symptoms, (aggressive behavior: *r*(50)=.28, *p*=.046 and rule-breaking behavior: *r*(50)=.34, *p*=.013) controlling for ADHD symptoms (Figure 2). CT and ADHD did not correlate when controlling for ODD/CD symptoms, *r*(50)=-.087, *p*=.54.

Regarding previous reports on a developmental delay of cortical thinning for ADHD (Shaw et al., 2007), we explored correlations of age and CT of the right rostral middle frontal cluster resulting for ADHD+ODD/CD versus ADHD-only (Figure 2) separately for the groups. Results showed cortical thinning in the ADHD-only, *r*(34)=-.61, *p*<.001, and TD, *r*(28)=-.44, *p*=.016, but not the ADHD+ODD/CD group, *r*(24)=-.11, *p*=.59. Fisher r-to-z transformation revealed a significant difference only between the correlation coefficients of the ADHD+ODD/CD and ADHD-only groups (*z*=-2.2, *p*=.028) but not of the ADHD+ODD/CD and TD (*z*=-1.25, *p*=.21) or ADHD-only and TD (*z*=-.93, *p*=.3) groups.

## Discussion

In accordance with two recent reports (Noordermeer et al., 2017; Stevens & Haney-Caron, 2012) this study is one of the first to provide knowledge to disentangle the related conditions ADHD-only and ADHD+ODD/CD. Going beyond previous studies, we analyzed several morphological markers, i.e. subcortical and cortical volume as well as CT and SA by contrasting a matched sample of ADHD-only with ADHD+ODD/CD. We found similar and differential alterations for the two patient groups. Similarly, both groups had reductions in total GM volume, right CV, and total SA. The ADHD+ODD/CD group had a reduced bilateral cerebellum while the ADHD-only group only had reductions in the right cerebellum. Differentially, ADHD+ODD/CD diverged from both TD and ADHD-only in a rostral middle frontal cluster. The increased thickness was related to ODD/CD symptoms under control of ADHD symptomatology. Further, we observed no thinning across adolescence in this cluster in the ADHD+ODD/CD, but not in the other groups. No other alterations for CV or SA survived stringent thresholding and ICV correction (Greve & Fischl, 2018).

In line with our hypothesis for global measures, both diagnostic groups had reductions in total GM volume with 4% for ADHD-only and 5% for ADHD+ODD/CD. Both groups also had reductions in right cortical GM volume (4% for ADHD-only and 5% for ADHD+ODD/CD), and total SA (4-5% for ADHD and 6% for ADHD+ODD/CD). This study adds evidence that for ADHD+ODD/CD total and cortical GM volume reductions are in a similar range as for ADHD-only (see Ambrosino et al., 2017 and Greven et al., 2015 for samples including comorbidity; see Noordermeer et al., 2017 for ODD). For SA, we extend previous findings that did not exclude ODD comorbidity (Wolosin et al., 2009) in showing that in both pure and comorbid ADHD similar reductions are evident in total SA.

Regarding alterations in CT ADHD+ODD/CD had a 6 to 7% thicker cortex than both ADHD-only and TD in a rostral middle frontal cluster. The region is in line with neuropsychological impairments in inhibition, a cognitive function that is dependent on the (inferior) frontal cortex and has been shown for both ADHD and with worse neuropsychological problems in ODD/CD (Loeber et al., 2000). The increased thickness in this cluster seems to be related to the ODD/CD and not ADHD component, since it correlated with ODD/CD symptoms when controlling for ADHD symptoms. Our results are in contrast to a study that found no CT alterations in ADHD with comorbid ODD (Noordermeer et al., 2017). A possible reason is that our sample was younger (mean age 16 versus 13, age range 7-29 versus 10-17). Two similar aged ADHD+ODD/CD studies found an increase in about the same location in the right prefrontal cortex in volume (that closely relates to thickness, Garrett et al., 2008; Semrud-Clikeman et al., 2014). The increased thickness in our cluster might also be related to a slower thinning in the ADHD+ODD/CD group since thickness did not decrease across adolescence in ADHD+ODD/CD. In contrast, for the TD and ADHD-only group a thinning was observed – as would be expected for typical adolescent development. Interestingly, a delay of attaining peak CT in a similar middle frontal region before adolescence was reported for ADHD at age 10.5 versus age 7.5 (Shaw et al., 2007). Almost half of the sample of the Shaw et al. (2007) study (42%) suffered from ODD/CD comorbidity. Probably, specifically the added burden with ODD/CD might relate to a delayed prefrontal cortical maturation in thickness. Future research could follow cortical maturation of ADHD with and without comorbid ODD/CD longitudinally to test whether the supposed later thinning specifically for ADHD with comorbid ODD/CD can be verified.

Partly as expected, we found reductions in the right cerebellum in both patient groups (6-8%), while in the left cerebellum only ADHD+ODD/CD differed from TD by 6%. Our results hint at the cerebellum reduction observed in ADHD (for a metaanalysis see Valera et al., 2007) is shared or even pronounced for ODD/CD comorbidity. However, this result did not survive FDR correction and therefore has to be dealt with caution.

We found no alterations for neither patient group for subcortical volumes similarly to two other adolescent studies (Ambrosino et al., 2017; Noordermeer et al., 2017). Sample age seems to be important as effect sizes for subcortical reductions were shown to be larger in child samples (see megaanalysis by Hoogman et al., 2017) and two meta-analyses showed that basal ganglia reductions tended to decrease with increasing age (Frodl & Skokauskas, 2012; Nakao et al., 2011).

We found no evidence of local alterations in CV and SA that survived robustness analyses. About only three previous studies report local SA alterations from which two assessed child samples and all three included participants with ODD comorbidity (Dirlikov et al., 2015; Silk et al., 2016; Wolosin et al., 2009). In line with these reports, for children but not adults with ADHD SA reductions have been shown (Hoekzema et al., 2012). Differences to previous studies demonstrating frontal volume alterations might also be related to sample age and inclusion of comorbidity in the ADHD sample (e.g. Ambrosino et al., 2017).

Overall, comparisons to previous reports remain difficult because - as was reported by a recent review (Noordermeer et al., 2016) - several previous studies using a whole brain approach did not control for multiple comparisons, nor applied cluster thresholding.

Limiting generalization of present findings, we assessed boys only; replication is warranted in mixed samples. Due to the small size of the ADHD+ODD/CD-group, we could not differentiate between ODD and CD alterations. Future studies are needed, that disentangle the specific contribution of ODD versus CD (Noordermeer et al., 2016; Noordermeer et al., 2017) and systematically compare ADHD-only, ADHD with comorbid ODD or with comorbid CD and the “pure forms” of ODD and CD. Due to the cross-sectional nature of our study, conclusions about a possible delayed cortical thinning in ADHD+ODD/CD remain to be verified by longitudinal studies.

### Conclusion

We demonstrate the necessity to carefully differentiate between ADHD and ADHD+ODD/CD since we found similarities (reduced volume of the total GM, SA, and the cerebellum) but also differences in brain morphology between the disorders (bilateral instead only right cerebellum reduced, increase in rostral middle frontal thickness for ADHD+ODD/CD). Specifically the increased rostral middle frontal thickness hints at a specific developmental brain alteration in ADHD+ODD/CD, probably related to a delayed cortical thinning. Implications for treatment could be to specifically focus on the (adolescent) boys with a “double burden” of ADHD+ODD/CD that seem to be even more affected than boys with ADHD-only.

## Acknowledgement

This research was supported by Bundesministerium für Bildung und Forschung, Grant Numbers:01EV0711, Aerial 01EE1406B; Deutsche Forschungsgemeinschaft, Grant Numbers: SFB 940/1, SFB 940/2, VE 892/2-1; Faculty of Medicine at the Technische Universität Dresden, MeDDrive Grant.

