## Supplement S1 for "Altered brain morphology in boys with attention-deficit/hyperactivity disorder with and without comorbid conduct disorder/oppositional defiant disorder"

**Supplement 1: Supplementary Methods**

**Procedure**

Participants completed two visits with 1–4 weeks in between. During the first visit, we assessed the WISC-IV and the M.I.N.I.-Kid. At the second visit, participants completed the MRI session. The T1-weighted sequence took place in between other two functional scans (an emotional attention paradigm (Vetter et al., 2018) and a probabilistic reversal learning paradigm, (Javadi, Schmidt, & Smolka, 2014). Participants and their parents filled in questionnaires in between these two visits.

**Image acquisition and processing**

3D T1-weighted magnetization-prepared rapid gradient echo (MPRAGE) image data sets were acquired (TR = 1900ms, TE = 2.26ms, FOV = 256x256mm, 176 slices, 1x1x1mm voxel size, flip angle = 9°) using a 3T whole-body MR tomograph (Magnetom TRIO, Siemens, Erlangen, Germany) equipped with a 12-channel head coil. Scan acquisition took 6 min.

FreeSurfer performs volumetric segmentation of global brain measures, the cerebellum, deep GM volumetric structures and subcortical WM (volume-based stream) and three-dimensional cortical reconstruction and parcellation (surface-based stream). In short, the processing includes removal of non-brain tissue using a hybrid watershed/surface deformation procedure (Ségonne et al., 2004), automated Talairach transformation and segmentation of the subcortical WM and deep GM), intensity normalization (Sled, Zijdenbos, & Evans, 1998), tessellation of the gray matter white matter boundary, automated topology correction (B. Fischl, Liu, & Dale, 2001; Ségonne, Pacheco, & Fischl, 2007), and surface deformation following intensity gradients to optimally place the gray/white and gray/cerebrospinal fluid borders at the location where the greatest shift in intensity defines the transition to the other tissue class (Dale, Fischl, & Sereno, 1999; Dale & Sereno, 1993; Bruce Fischl & Dale, 2000).

Volume-based ROIs were obtained from FreeSurfer’s *aseg.stats files. Cortical measures (CV, CT, SA) were analyzed and corrected for multiple comparisons using the command-line group analysis stream in FreeSurfer including assembling the data and general linear modeling.

Subcortical GM volume was calculated manually by adding bilateral volumes of the thalamus, caudate, putamen, pallidum, hippocampus, amygdala and accumbens.

**Cortical reconstruction**

Cortical measures of interest outputted by the surface-based stream were cortical volume (CV), white surface area (SA) and cortical thickness (CT) of the (whole brain/vertex-wise approach). SA was calculated as the area of the triangles at the pial surface touching each vertex in the tessellated surface representation. CT was calculated as the mean distance between vertices of a corrected, triangulated estimated GM/WM surface and GM/CSF (pial) surface (Bruce Fischl & Dale, 2000). CV was calculated as each vertex’s CT multiplied by SA at the midway point between the white and pial surfaces.

**Quality control**

To ensure high data quality we followed a workflow, which included screening T1-weighted images immediately after the scanning session and rescanning in case of bad data quality. We coded all T1-weighted scans in order to ensure rater blindness to participants’ diagnosis. To ensure high data quality we followed a workflow which included screening T1-weighted images immediately after the scanning session, rescanning the participant in case of bad data quality (Backhausen et al., 2016). To prevent movement we used cushions stabilizing the head and participants could undertake a Mock scanner session to get used to the MRI environment and practice laying still. The FreeSurfer pipeline requires that data is checked individually at multiple points. As volume-based and surface-based measures are sensitive to movement and other acquisition artifacts, a thorough visual quality control (QC) ensures robustness and reliability of results (Ducharme et al., 2016).

We applied detailed visual quality control for 1) raw T1-weighted images and 2) processed images after completing the FreeSurfer processing pipeline (recon-all) in line with previously published guidelines of our group (Backhausen et al., 2016). Raw T1-weighted images were visually rated on the four steps “Image sharpness”, “Ringing”, “Contrast to noise ratio (GM/WM)” and “Contrast to noise ratio (subcortical structures)” and assigned to one of three categories (pass, warn or fail). Images falling into the “fail” category were excluded from further analyses and FreeSurfer processed images of the other two categories underwent visual QC taking into account the deformation of the 3D brain anatomy, removal of non-brain tissue, i.e. “skullstrip”, plausibility of subcortical/cortical structure borders and absence of GM misclassification as WM (Backhausen et al., 2016).

After visual quality control, we increased the watershed parameter for six participants (two ADHDpure group, ADHDcomo group, three TD group) in order to improve skull stripping. No other manual correction of FreeSurfer segmentations was performed.

***Statistical analysis***

For volume-based ROIs, we performed robustness analyses to assess the stability of findings. First, we added ICV as a covariate to control for differences in head size. Second, we applied a false discovery rate (FDR) Benjamini–Hochberg multiple comparisons correction of *p*<.05. All volume-based ROIs as well as the global measures CV, mean CT and total SA were analyzed separately by hemisphere.
