## Supplementary material for "Altered brain morphology in boys with attention-deficit/hyperactivity disorder with and without comorbid conduct disorder/oppositional defiant disorder": Table S1

Global volume and volume-based ROIs: results of group comparisons.

| **Measure** | L/R | Volume (mm^3^) | | | *F* | *p* | *Post-Hoc Group Comparisons* |
| --- | --- | --- | --- | --- | --- | --- | --- |
|  |  | **ADHD-only**  ***n*=36**  *M (SD*) | **ADHD+ ODD/CD**  ***n*=26**  *M (SD*) | **TD**  ***n*=30**  *M (SD*) |  |  |  |
| **Global measures** | |  |  |  |  |  |  |
| Intracranical volume |  | 1,534,105 (127,186) | 1,539,709 (89,903) | 1,615,853 (124,760) | 4.67 | .012 | ADHD-only<TD^2^ , ADHD+ODD/CD<TD^2^ |
| Total GM volume |  | 803,197 (48,484) | 795,511 (55,391) | 836,658 (57,473) | 4.91 | .01 | ADHD-only<TD^2^, |
| Cortical GM volume | R | 295,765 (20,556) | 293,752 (23,719) | 307,879 (24,902) | 3.28 | .042 | ADHD-only<TD^2^, |
|  | L | 293,196 (20,662) | 290,981 (25,825) | 304,93 (26,149) | 2.87 | .062 |  |
| Total SA | R | 91,102 (6,824) | 89,694 (7,234) | 94,851 (6,179) | 4.52 | .014 | ADHD-only<TD^1^, ADHD+ODD/CD<TD^1^ |
|  | L | 90,051 (6,714) | 89,148 (7,066) | 94,175 (6,447) | 4.66 | .012 | ADHD-only<TD^1^, ADHD+ODD/CD<TD^1^ |
| **Volume-based ROIs** |  |  |  |  |  |  |  |
| Cerebellum (GM) | R | 66,392 (5,012) | 65,133 (4,032) | 70,203 (5,407) | 5.03 | .009 | ADHD-only<TD^1^, ADHD+ODD/CD <TD^1^ |
|  | L | 64,877 (4,957) | 64,225 (3,843) | 68,284 (5,393) | 3.16 | .047 | ADHD+ODD/CD<TD^2^ |

*Note*. *F* and *p* values of the volume-based ROIs refer to the main effect of group in the MANCOVA including ICV as covariate. Post-hoc results are group differences using t-tests for global measures and ANCOVAS with two groups with ICV as covariate for volume-based ROIs. *^1^p*<.01, *^2^p*<.05. ADHD-only= attention-deficit hyperactivity disorder group, ADHD+ODD/CD = comorbid attention-deficit hyperactivity disorder and conduct disorder/oppositional defiant disorder group, TD=typically developing group, GM = gray matter, SA = surface area
