## Supplementary material for "Altered brain morphology in boys with attention-deficit/hyperactivity disorder with and without comorbid conduct disorder/oppositional defiant disorder": Table S2

*Group comparisons applying liberal CFT thresholds*

*Differences in cortical thickness; CFT p < .05*

| Cortical region | L/R | MNI coordinates | | | Cluster size (mm^2^) | Number of vertices | Clusterwise p-value (CWP) | Post-Hoc Group Comparisons |
| --- | --- | --- | --- | --- | --- | --- | --- | --- |
|  |  | X | Y | Z |  |  |  |  |
| **ADHD-only vs ADHD+ODD/CD vs TD** | | | | | | | |  |
| Rostral middle frontal | R | 44 | 23 | 29 | 2436 | 4189 | .003 |  |
| **ADHD+ODD/CD vs TD** | | | | | | | |  |
| Rostral middle frontal | L | -23 | 50 | 19 | 898 | 1268 | .553 | ADHD+ODD/CD>TD |
|  | R | 44 | 23 | 30 | 3280 | 5510 | <.001 | ADHD+ODD/CD>TD |
| Inferior parietal | L | -34 | -61 | 39 | 447 | 926 | .1 | ADHD+ODD/CD>TD |
|  | R | -38 | -70 | 45 | 807 | 1654 | .681 | ADHD+ODD/CD>TD |
| Superior frontal | L | -12 | -5 | 65 | 749 | 1430 | .781 | ADHD+ODD/CD>TD |
| **ADHD-only vs ADHD+ODD/CD** | | | | | | | |  |
| Rostral middle frontal | L | -34 | 33 | 30 | 605 | 905 | .947 | ADHD-only< ADHD+ODD/CD |
|  | R | 36 | 39 | 10 | 3137 | 5104 | <.001 | ADHD-only< ADHD+ODD/CD |
| Frontal pole | R | 11 | 58 | -14 | 600 | 810 | .948 | ADHD-only< ADHD+ODD/CD |
| Superior frontal | R | 10 | 46 | 21 | 488 | 876 | .995 | ADHD-only< ADHD+ODD/CD |
| Inferior parietal | R | 41 | -66 | 37 | 445 | 814 | .999 | ADHD-only< ADHD+ODD/CD |

*Note:* Smoothing kernel = 15mm FWHM, cluster-forming threshold p < .05, Monte Carlo Z simulation with 10000 iterations; CFT = cluster-forming threshold, CWP = clusterwise p-value. ADHD-only= attention-deficit hyperactivity disorder group, ADHD+ODD/CD = comorbid attention-deficit hyperactivity disorder and conduct disorder/oppositional defiant disorder group, TD=typically developing group.

*Differences in cortical volume; CFT p <* .05

| Cortical region | L/R | MNI coordinates | | | Cluster size (mm^2^) | Number of vertices | Clusterwise p-value (CWP) | Post-Hoc Group Comparisons |
| --- | --- | --- | --- | --- | --- | --- | --- | --- |
|  |  | X | Y | Z |  |  |  |  |
| **ADHD-only vs ADHD+ODD/CD vs TD** | | | | | | | |  |
| Rostral middle frontal | R | 39 | 52 | -4 | 1735 | 2450 | .105 |  |
| Fusiform | R | 40 | -63 | -17 | 769 | 1274 | .913 |  |
|  | R | 34 | -11 | -34 | 567 | 1060 | .995 |  |
| Superior parietal | L | 22 | -82 | 15 | 554 | 858 | .999 |  |
| **ADHD-only vs TD** | | | | | | | |  |
| Rostral middle frontal | R | 39 | 51 | -5 | 2284 | 3231 | .019 | ADHD-only<TD |
| Precuneus | R | 24 | -61 | 17 | 765 | 1477 | .916 | ADHD-only<TD |
| Fusiform | R | 34 | -10 | -35 | 698 | 1381 | .959 | ADHD-only<TD |
| Pars orbitalis | L | -38 | 38 | -11 | 679 | 1188 | .98 | ADHD-only<TD |
| **ADHD+ODD/CD vs TD** | | | | | | | |  |
| Fusiform | R | 34 | -12 | -33 | 645 | 1220 | .981 | ADHD+ODD/CD<TD |
|  | R | 41 | -53 | -13 | 636 | 1108 | .983 | ADHD+ODD/CD<TD |
| **ADHD-only vs ADHD+ODD/CD** | | | | | | | |  |
| Frontal pole | R | 13 | 58 | -10 | 861 | 1180 | .826 | ADHD-only< ADHD+ODD/CD |
| Middle temporal | R | 47 | -62 | 5 | 778 | 1410 | .905 | ADHD-only> ADHD+ODD/CD |
| Fusiform | R | 40 | -65 | -16 | 771 | 1158 | .912 | ADHD-only> ADHD+ODD/CD |
| Superior parietal | L | -23 | -81 | 15 | 736 | 1142 | .958 | ADHD-only< ADHD+ODD/CD |
| Lingual | R | 7 | -90 | -11 | 724 | 790 | .944 | ADHD-only> ADHD+ODD/CD |
|  | L | -13 | -77 | -7 | 619 | 648 | .993 | ADHD-only> ADHD+ODD/CD |

*Note:* Smoothing kernel = 15mm FWHM, cluster-forming threshold p < .05, Monte Carlo Z simulation with 10000 iterations; CFT = cluster-forming threshold, CWP = clusterwise p-value. ADHD-only= attention-deficit hyperactivity disorder group, ADHD+ODD/CD = comorbid attention-deficit hyperactivity disorder and conduct disorder/oppositional defiant disorder group, TD=typically developing group.

*Differences in surface area; CFT p < .05*

| Cortical region | L/R | MNI coordinates | | | Cluster size (mm^2^) | Number of vertices | Clusterwise p-value (CWP) | Post-Hoc Group Comparisons |
| --- | --- | --- | --- | --- | --- | --- | --- | --- |
|  |  | X | Y | Z |  |  |  |  |
| **ADHD-only vs ADHD+ODD/CD vs TD** | | | | | | | |  |
| Middle temporal | R | 48 | -61 | 4 | 1204 | 2173 | .32 |  |
| Rostral middle frontal | R | 38 | 51 | -3 | 906 | 1300 | .668 |  |
| Fusiform | R | 34 | -11 | -34 | 609 | 1152 | .975 |  |
|  | R | 41 | -60 | -17 | 494 | 871 | .998 |  |
| Superior parietal | L | -22 | -84 | 14 | 564 | 881 | .99 |  |
| Pars orbitalis | L | -38 | 40 | -12 | 537 | 919 | .995 |  |
| **ADHD-only vs TD** | | | | | | | |  |
| Rostral middle frontal | R | 39 | 52 | -5 | 1537 | 2210 | .115 | ADHD-only<TD |
| Fusiform | R | 35 | -10 | -36 | 635 | 1199 | .963 | ADHD-only<TD |
| Pars orbitalis | L | -39 | 39 | -12 | 683 | 1198 | .936 | ADHD-only<TD |
| Superior parietal | R | 31 | -72 | 19 | 657 | 1340 | .95 | ADHD-only<TD |
|  | L | -22 | -85 | 14 | 591 | 945 | .984 | ADHD-only<TD |
|  | L | -33 | -44 | 54 | 509 | 1056 | .998 | ADHD-only<TD |
| **ADHD+ODD/CD vs TD** | | | | | | | |  |
| Rostral middle frontal | R | 39 | 52 | -2 | 770 | 1065 | .844 | ADHD+ODD/CD<TD |
| Inferior parietal | R | 34 | -71 | 21 | 731 | 1252 | .889 | ADHD+ODD/CD<TD |
| Fusiform | R | 33 | -11 | -33 | 691 | 1319 | .925 | ADHD+ODD/CD<TD |
| Middle temporal | R | 48 | -60 | 3 | 640 | 1158 | .960 | ADHD+ODD/CD<TD |
| Pars orbitalis | L | -43 | 42 | -10 | 772 | 1209 | .856 | ADHD+ODD/CD<TD |
| Rostral anterior cingulate | L | -12 | 41 | 6 | 696 | 1257 | .926 | ADHD+ODD/CD<TD |
| **ADHD-only vs ADHD+ODD/CD** | | | | | | | |  |
| Middle temporal | R | 48 | -61 | 4 | 1192 | 2053 | .329 | ADHD-only> ADHD+ODD/CD |
| Fusiform | R | 41 | -61 | -17 | 614 | 914 | .973 | ADHD-only> ADHD+ODD/CD |
| Lingual | R | 9 | -87 | -12 | 475 | 481 | .999 | ADHD-only> ADHD+ODD/CD |
|  | L | -11 | -77 | -6 | 866 | 973 | .738 |  |
| Superior parietal | L | -22 | -85 | 24 | 632 | 976 | .967 | ADHD-only< ADHD+ODD/CD |
| Lateral occipital | L | -41 | -76 | 3 | 563 | 909 | .991 | ADHD-only> ADHD+ODD/CD |

*Note:* Smoothing kernel = 15mm FWHM, cluster-forming threshold p < .05, Monte Carlo Z simulation with 10000 iterations; CFT = cluster-forming threshold, CWP = clusterwise p-value. ADHD-only= attention-deficit hyperactivity disorder group, ADHD+ODD/CD = comorbid attention-deficit hyperactivity disorder and conduct disorder/oppositional defiant disorder group, TD=typically developing group.
