## Supplementary material for "Altered brain morphology in boys with attention-deficit/hyperactivity disorder with and without comorbid conduct disorder/oppositional defiant disorder": Table S3

*Group comparisons applying stringent CFT thresholds*

*Differences in cortical volume; CFT p <* .005

| Cortical region | L/R | MNI coordinates | | | Cluster size (mm^2^) | Number of vertices | Clusterwise p-value (CWP) | Post-Hoc Group Comparisons |
| --- | --- | --- | --- | --- | --- | --- | --- | --- |
|  |  | X | Y | Z |  |  |  |  |
| **ADHD-only vs ADHD+ODD/CD vs TD** | | | | | | | |  |
| Rostral middle frontal | R | 39 | 52 | -4 | -418 | 26 | 0.998 |  |
| Middle temporal | R | 47 | -62 | 5 | 47 | 98 | 0.993 |  |
| Precuneus | R | 24 | -60 | 18 | 27 | 52 | 0.997 |  |
| Fusiform | R | 34 | -11 | -34 | 172 | 283 | 0.833 |  |
| Pars orbitalis | L | -38 | 38 | -11 | 1 | 2 | 0.998 |  |
| Lateral occipital | L | -43 | -78 | 3 | 43 | 66 | 0.988 |  |
| Medial orbito frontal | L | -10 | 42 | -12 | 12 | 22 | 0.996 |  |
| Superior parietal | L | -23 | 82 | 15 | 133 | 217 | 0.89 |  |
| **ADHD-only vs TD** | | | | | | | |  |
| Rostral middle frontal | R | 39 | 51 | -5 | 366 | 527 | 0.344 | ADHD-only<TD |
| Precuneus | R | 24 | -61 | 17 | 127 | 255 | 0.923 | ADHD-only<TD |
| Fusiform | R | 34 | -10 | -35 | 166 | 273 | 0.848 | ADHD-only<TD |
| Medial orbito frontal | L | -10 | 42 | -12 | 126 | 204 | 0.902 | ADHD-only<TD |
| Pars orbitalis | L | -38 | 38 | -11 | 116 | 193 | 0.92 | ADHD-only<TD |
| **ADHD+ODD/CD vs TD** | | | | | | | |  |
| Fusiform | R | 34 | -12 | -33 | 240 | 417 | 0.663 | ADHD+ODD/CD<TD |
|  | R | 41 | -53 | -13 | 155 | 268 | 0.871 | ADHD+ODD/CD<TD |
| **ADHD-only vs ADHD+ODD/CD** | | | | | | | |  |
| Frontal pole | R | 13 | 58 | -10 | 67 | 72 | 0.985 | ADHD-only< ADHD+ODD/CD |
| Middle temporal | R | 47 | -62 | 5 | 147 | 309 | 0.883 | ADHD-only> ADHD+ODD/CD |
| Lateral occipital | L | -42 | -77 | 3 | 148 | 232 | 0.862 | ADHD-only> ADHD+ODD/CD |
| Superior parietal | L | -23 | -81 | 15 | 278 | 444 | 0.53 | ADHD-only< ADHD+ODD/CD |

*Note:* Smoothing kernel = 15mm FWHM, cluster-forming threshold p < .005, Monte Carlo Z simulation with 10000 iterations; CFT = cluster-forming threshold, CWP = clusterwise p-value. ADHD-only= attention-deficit hyperactivity disorder group, ADHD+ODD/CD = comorbid attention-deficit hyperactivity disorder and conduct disorder/oppositional defiant disorder group, TD=typically developing group.

*Differences in surface area; CFT p <* .001

| Cortical region | L/R | MNI coordinates | | | Cluster size (mm^2^) | Number of vertices | Clusterwise p-value (CWP) | Post-Hoc Group Comparisons |
| --- | --- | --- | --- | --- | --- | --- | --- | --- |
|  |  | X | Y | Z |  |  |  |  |
| **ADHD-only vs ADHD+ODD/CD vs TD** | | | | | | | |  |
| Middle temporal | R | 48 | -61 | 4 | 68 | 130 | 0.589 |  |
| Fusiform | R | 34 | -11 | -34 | 15 | 22 | 0.824 |  |
| Lateral occipital | L | -42 | -76 | 4 | 56 | 85 | 0.647 |  |
| **ADHD-only vs TD** | | | | | | | |  |
| Fusiform | R | 35 | -10 | -36 | 8 | 11 | 0.849 | ADHD-only<TD |
| **ADHD+ODD/CD vs TD** | | | | | | | |  |
| Rostral middle frontal | R | 39 | 52 | -2 | 39 | 54 | 0.720 | ADHD+ODD/CD<TD |
| Fusiform | R | 33 | -11 | -33 | 16 | 24 | 0.822 | ADHD+ODD/CD<TD |
| **ADHD-only vs ADHD+ODD/CD** | | | | | | | |  |
| Middle temporal | R | 48 | -61 | 4 | 138 | 270 | 0.323 | ADHD-only> ADHD+ODD/CD |
| Lateral occipital | L | -41 | -76 | 3 | 103 | 160 | 0.447 | ADHD-only> ADHD+ODD/CD |

*Note:* Smoothing kernel = 15mm FWHM, cluster-forming threshold p < .001, Monte Carlo Z simulation with 10000 iterations; CFT = cluster-forming threshold, CWP = clusterwise p-value. ADHD-only= attention-deficit hyperactivity disorder group, ADHD+ODD/CD = comorbid attention-deficit hyperactivity disorder and conduct disorder/oppositional defiant disorder group, TD=typically developing group.
